# Genome restructuring and adaptation in Arctic marine bacteria

**DOI:** 10.1101/2025.02.13.637858

**Authors:** Mike C. Sadler, Matthias Wietz, Sayaka Mino, Robert M. Morris

## Abstract

Arctic marine bacteria experience seasonal changes in temperature, salinity, and light caused by the formation and melting of sea-ice. Time-series studies have identified spatial and temporal patterns in microbial communities in the Arctic and environmental sequencing has provided insights into the genetic potential of key taxa. We cultured and sequenced the complete genomes of 34 Arctic marine bacteria to identify patterns of gene gain, loss, and rearrangement that structure genomes and underlie adaptations to Arctic conditions. We found that the most abundant lineage in the Arctic (SAR11) is comprised of diverse species and subspecies, each encoding 50-150 unique genes. Half of the SAR11 genomes (8/16) harbor a genomic island with the potential to enhance survival in the Arctic by utilizing the osmoprotectant and potential methyl donor glycine betaine. We also cultured and sequenced four species from a new family of *Pseudomonadales*, four subspecies of *Pseudothioglobus* (SUP05), a genus of high GC *Puniceispirillaes* (SAR116), and a family of low GC SAR116. Time-series data indicate that this collection represents up to 60% of the marine bacterial community in Arctic waters at peak abundance. Their complete genomes provide insights into the evolutionary processes that underlie diversity and adaptation to the Arctic Ocean.

## Introduction

The oceans are dominated by relatively few but highly diverse lineages of marine bacteria [1, 2]. Representatives from some of the most abundant lineages have been cultured, including *Pelagibacterales* (SAR11) [3, 4], *Puniceispirillales* (SAR116) [5], and *Thioglobaceae* (SUP05) [6, 7], but there is significant diversity in natural populations. Progress studying diversity has been made using metagenome assembled genomes (MAGs) [8–10] and single-cell amplified genomes (SAGs) [11–13]. The challenge is that MAGs and SAGs are typically incomplete and do not provide a full picture of the genetic variation within a highly diverse population of species and strains [14]. Advances in cultivation have resulted in the recovery of abundant and novel marine taxa [15], while modern sequencing techniques produce highly accurate closed bacterial genomes [16, 17]. Combined, these approaches can provide more complete information about the genetic variation within populations.

The Arctic Ocean is a highly dynamic marine system characterized by strong spatial and temporal variation, including sea ice cover, extended darkness, stratification, and freshwater input [18–20]. Marine microbes have evolved specific adaptations to survive in these extremes, such as the ability to use the compatible solute glycine betaine [21], which can enhance osmoregulation and survival in sea ice [22, 23] or can serve as a methyl donor and source of glycine [24]. Some Arctic bacteria have been isolated in culture, including *Colwellia* and *Polaribacter* [25–27]. However, cultivation independent methods suggest that current Arctic culture collections represent a relatively small fraction of the community [28]. Understanding the genetic variation within populations of Arctic bacteria is particularly important because the Arctic is warming at four times the global rate [29], rapidly losing sea ice [30], and experiencing stronger Atlantic water intrusion [31]. These changes have the potential to impact microbial diversity [32].

We conducted a population genomics study to gain insights into the genomic diversity within Arctic bacterial populations. We focused on highly abundant and diverse taxa with few cultured and sequenced representatives, such as SAR11 [33, 34]. SAR11 account for approximately 25% of bacteria in the Arctic Ocean [35–37] and are also present in sea ice [38]. Like many other marine bacterial lineages, the SAR11 clade has been classified into subclades based on 16S rRNA and internal transcribed spacer (ITS) sequence analysis [39, 40]. These classifications are often used to identify patterns of diversity, including in the Arctic [41–43]. SAR11 genomes are among the smallest for free living bacteria [44] and have a high proportion of core genes [40], with most unique genes co-located in a ∼50 Kb hypervariable region (HVR) termed HVR2 [45]. The evolutionary mechanisms that maintain diversity in HVRs are poorly understood, though homologous recombination is widespread in SAR11 [46] and has been proposed as a driver of diversity in marine bacteria [47]. These features have hindered efforts to resolve genetic variation and recover unique genes within populations of SAR11 [14].

This study used a cultivation-based approach to identify patterns of gene gain, loss, and rearrangement in Arctic populations of SAR11 and other previously uncultured strains, species, genera, and families of Arctic bacteria, named herein. Rationale for names assigned to previously uncultured families and genera are summarized below and detailed in the protologue. Briefly, *Candidatus* Njordibacter, the genus name derived from “Njord” (the Norse god of wind and seas), and the family *Njordibacteraceae* to encompass this genus. *Candidatus* Levibacter, the genus name derived from the Latin “levis” (light weight), and the family *Levibacteraceae* to encompass this low GC genus of *Puniceispirillales*. *Candidatus* Ponderosibacter, the genus name derived from the Latin “ponderosus” (heavy weight), referring to this genus of the high GC *Puniceispirillales*. *Candidatus* Marifrigibacter, the genus name derived from the Latin “mare” (of the sea) and “frigus” (cold). All 34 new species names correspond to their cultivation ID (e.g. sp. uisw_002).

## Materials and Methods

### Sample collection

Seawater samples were collected in the Arctic Ocean aboard the RV Kronprins Haakon from May 18 – 21 2023, through ∼2 m thick sea ice. Samples for cultivation (50 mL) were collected at 81.04° N, 10.62° E, from 25 m below the ice/water interface. A 1 mL seawater sample was amended to 10% (v/v) glycerol, flash frozen in liquid nitrogen, and stored at -80 °C until used for high-throughput dilution to extinction cultivation.

Seawater for culture media (10 L) was collected at 80.96° N, 9.66° E from 1 m below the ice/water interface into acid washed and miliQ rinsed cubitainers. Seawater was then filter-sterilized using a tangential flow filtration (TFF) system equipped with a 30 kDa Pellicon XL Polyethersulfone Biomax filter (MilliporeSigma, Burlington, MA). The resulting media was collected in 1 L acid washed and autoclaved polycarbonate bottles, incubated for 2 months at 4 °C, and checked for bacterial growth to ensure sterility prior to use.

### Bacteria cultivation

Cultures were obtained by high-throughput dilution to extinction cultivation using cryopreserved Arctic seawater as previously described [48]. Briefly, cryopreserved cells were diluted to ∼33 cells per mL in 30 kDa filter sterilized seawater media. 1.5 mL was then added to each well of an acid washed and sterile 96 well Teflon plate, incubated at 4 °C, and monitored for growth once a week for 13 weeks using a Guava easyCyte flow cytometer (Cytek, Fremont, CA). Wells that were positive for growth (>2 x 10^4^ cells per mL) were identified by 16S rRNA sequence analysis when cell densities reached >10^5^ cells per mL and preserved in 10% (v/v) glycerol that was frozen and stored at -80 °C.

### Culture identification

DNA for PCR was extracted from 100 µL of cell culture and sequenced using a physical lysis procedure as previously described [49]. Briefly, potassium hydroxide – dithiothreitol was added and samples were subjected to 1 freeze thaw cycle. The pH was then adjusted to 8.0 using tris-HCl. DNA was purified using 2X volume:volume DNA mag beads (Sergi Lab Supplies, Seattle, WA) with two 80% ethanol washes, and eluted in 20 µL 10 mM tris-HCL. The 16S rRNA gene was then amplified using PCR with primers 27F_B (5’ AGRGTTYGATYMTGGCTCAG 3’) and 926R_B (5’ CCGYCAATTCMTTTRAGTTT 3’), and in some cases using a second semi-nested PCR reaction with primers 519F (5’ CAGCMGCCGCGGTAATWC 3’) and 926R_B. The following PCR conditions were used throughout: 38 cycles (94 °C 20 s, 55 °C 45 s, 72 °C 120 s). PCR products were sequenced by Genewiz (Genewiz, Seattle, WA). Cultures were putatively identified by aligning sequences with related sequences in the Silva database v138.1 [50, 51].

### Genome sequencing

Cultures selected for whole genome sequencing were revived from freezer stocks in 1 L acid washed and autoclaved polycarbonate bottles containing TFF sterilized Puget Sound seawater media. Cells were collected on 47 mm 0.2 µm pore size Isopore membrane filters (MilliporeSigma, Burlington, MA) when cultures reached maximum cell densities (between 10^5^ and 10^6^ cells per mL). High molecular weight DNA was extracted using the Autogen Quickgene DNA Tissue Kit (Autogen, Holliston, MA) following the extraction protocol for animal tissue with minor modifications as noted below. Filters were cut into small pieces using sterile forceps and scissors and placed in sterile 2 mL DNA LoBind tubes (Eppendorf, Hamburg, Germany) containing 200 µL of TE buffer. Filters were then frozen at -80 °C for 20 minutes and heated until thawed at 95 °C. All recommended extraction volumes were doubled, and DNA was eluted in 200 µL of molecular grade water. DNA was cleaned using 1X volume:volume DNA magnetic beads (Sergi Lab Supplies, Seattle WA) with two 80% ethanol washes, then eluted in 20 µL of molecular grade water. DNA was quantified using a Qubit dsDNA Quantitation High Sensitivity kit (Invitrogen, Waltham, MA) and sequenced using the Oxford Nanopore Technologies (ONT) R10.4.1 Flongle flow cells with the SQK-RAD114 rapid library prep kit (Oxford Nanopore Technologies, Oxford, United Kingdom). Bases were called with Dorado (github.com/nanoporetech/dorado), using the dna_r10.4.1_e8.2_400bps_hac@v4.2.0 model.

### Genome assembly and annotation

Bacterial genomes were assembled with Flye v2.9.1 [52] and polished with Medaka v1.7.2 (github.com/nanoporetech/medaka) using the usegalaxy web platform [53]. Genome annotation was performed by NCBI using the Prokaryotic Genome Annotation Pipeline v6.8 [54, 55]. The quality of ONT genomes was evaluated by comparing ONT-only genomes constructed with varying levels of coverage (9-500x) to ONT-Illumina hybrid genome assemblies obtained for two previously sequenced strains, SAR11 strain NP1 [56] and SUP05 strain EF3 [57]. Coverage for ONT-only genomes was varied by subsampling ONT reads (accessions: SRX26378910 & SRX22361185) with Rasusa v2.0.0 [58]. Hybrid genomes were created by polishing ONT genomes with Illumina reads (accessions: SRX26378911 & SRX23025519) using BWA-mem2 v2.2.1 [59] and Pilon v1.2.0 [60]. The quality of ONT-only genomes was determined by identifying the number of mismatches, indels, and excess CDSs relative to hybrid assemblies using Quast v5.2.0 [61] (**Fig. S1**). Only closed genomes with greater than 10 X coverage, corresponding to >99.9% accuracy, were used for further analyses.

### Spatial and temporal abundance

16S rRNA sequences of cultures were matched against 5,511 ASVs derived from the FRAM (FRontiers in Arctic marine Monitoring) Observatory in Fram Strait. Data originated from year-round autonomous sampling in the polar-influenced East Greenland Current [62] and the Atlantic-influenced West Spitsbergen Current [63] between 2016 – 2020, with approximately biweekly resolution. Using Genious Prime, we identified ASVs with 100% matches to Arctic isolates and plotted their abundances in Fram Strait over time to establish a broader spatiotemporal and ecological context.

### Phylogenomics and population genomics

Taxonomic classifications were assigned to sequenced genomes using the GTDB-Tk v2.3.2, with the de-novo workflow and reference database v214 [64]. New species and strains were given the prefix “uisw” followed by the cultivation number. Most genomes came from pure cultures (n=27). The suffix “_01” or “_02” was added to the cultivation number if two complete genomes were recovered from a mixed culture. The genomes of cultured isolates were analyzed using average amino acid identity (AAI) with ezaai v1.2.3 [65], percent of conserved proteins (POCP) with POCP v2.3.2 [66], and average nucleotide identity (ANI) with pyANI v0.2.12 [67]. Genome structure was evaluated through the visualization of linear co-similarity blocks using progressiveMauve v2015-02-25 [68]. Phylogenetic trees were constructed with MUSCLE v3.8.31 [69] and RAxML v8.2.11 with model GTRGAMMA [70] using the ETE3 v3.1.3 phylogenetic analysis pipeline [71]. Whole genome phylogenies were constructed using the bacterial_71 single-copy core gene collection in anvi’o v8 [72]. SAR11 gene cluster data was created using the anvi’o pangenomic workflow with the flags –use-ncbi-blast, –minbit 0.5, and –mcl-inflation 10, as previously described [73].

## Results

### Cultivation-based whole genome sequencing of Arctic marine bacteria

We cultured and sequenced the complete genomes of 34 Arctic marine bacteria to identify differences in genome structure and adaptations to Arctic conditions (**Fig. 1**). Cultures were selected from 106 bacteria obtained by high-throughput dilution to extinction cultivation and sequenced using the Oxford Nanopore Technologies (ONT) platform (**Table S1)**. All genomes are single circular contigs, range in size from 1.29 to 3.78 Mbp, and have GC contents between 29 and 52% (**Table S2**). They include representatives from well-known lineages, such as *Pelagibacter* (SAR11, n=16), *Pseudothioglobus* (SUP05, n=4), *Puniceispirillales* (SAR116, n=2), *Methylophilaceae* (OM43, n=2), and *Haliaceae* (OM60, n=1), as well as uncultured and understudied lineages, including a low GC family of SAR116 [74] and a novel family of *Pseudomonadales*. New genera and species of *Puniceispirillales*, *Pseudomonadales*, *Flavobacteriaceae*, and *Actinomycetes* were also cultured.

**Fig. 1.**
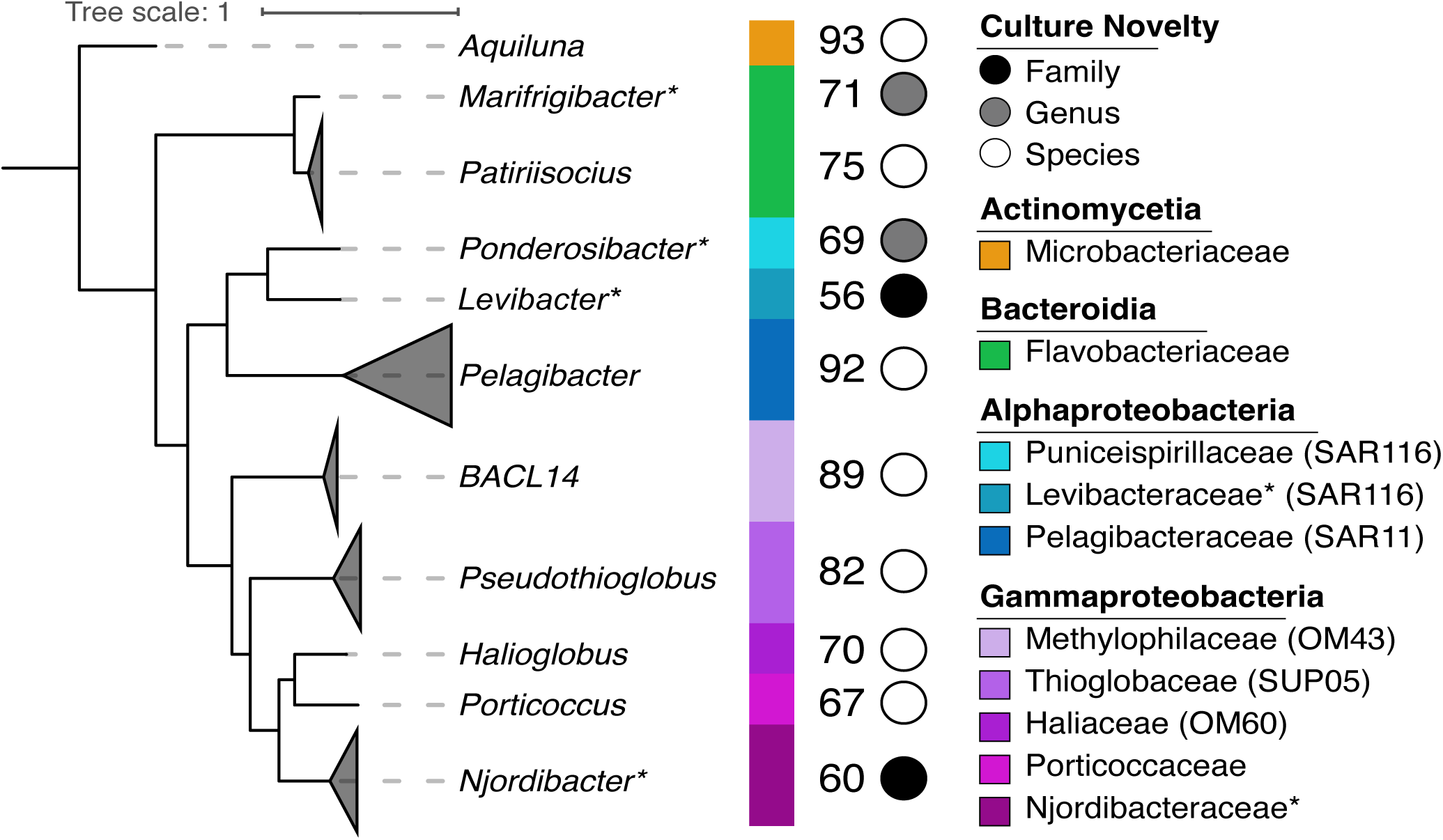
Diversity of bacteria isolated and sequenced from the Arctic Ocean. Phylogenomic analysis of Arctic marine bacteria sequenced in this study, with genus affiliation (italics), class and family taxonomic grouping (color bars), amino acid identity (AAI) to closest cultured relative (number), and culture novelty in circles (family, genus, and species). Wedges are proportional to the number of cultures sequenced for each lineage. Family and genus names were assigned using the GTDB-Tk placement tool, except for those proposed in this study, marked with an asterisk.

All isolates represent newly cultured species (**Fig. 1, Table S2**). The four SUP05 genomes have 95% ANI when compared to each other and less than 95% ANI when compared to previously cultured species. The SUP05 genomes code for key proteins needed to fix inorganic carbon using the Calvin-Benson-Bassham (CBB) cycle, including phosphoribulokinase (PRK) and the large and small subunits of form Ia Ribulose Bisphosphate Carboxylase (RuBisCO). *Levibacter* and *Ponderosibacter* belong to the SAR116 clade and have 56 and 69% AAI when compared to their nearest cultured relatives, indicating that they derive from a previously uncultured family and genus, respectively. The SAR116 clade forms high and low GC subclades with 50±7 and 31±1 GC content, respectively [74]. The *Levibacter* genome has a GC content of 31.4% and therefore represents the first low GC SAR116 culture. The *Porticoccus, Patiriisocius,* and *Halioglobus* genomes have low AAI values when compared to their nearest cultured relatives (67, 75, and 70%, respectively), indicating significant diversity in these lineages. The *Marifrigibacter* genome has a similarly low AAI value when compared to its nearest cultured relative (71%) and represents the first isolate of an uncultured genus of *Flavobacteriaceae*. Notably, the *Porticcocus* genome has 67% AAI and is 40% (1.4 Mbp) smaller than its nearest cultured relative.

### Spatial and temporal abundance of cultured taxa

Spatiotemporal abundance estimates of cultured Arctic lineages were determined by matching the 16S rRNA gene sequences from cultures to amplicon sequence variants (ASVs) from two four-year time-series datasets: one from the West Spitsbergen Current (WSC) and one from the East Greenland Current (EGC) [62, 63]. The 34 cultures matched 18 ASVs with 100% identity (**Fig. 2**), collectively comprising over 60% of relative abundance in the WSC and 50% of relative abundance in the EGC (**Fig. 2A**). The most abundant members in the culture collection included SAR11, SUP05, and *Njordibacter*, comprising 10, 4, and 4% of the community in the WSC, and 15, 8, and 6% of the community in the EGC at peak abundance, respectively (**Fig. 2B**). While four/116 SAR11, one/55 SUP05, and four/24 *Njordibacter* ASVs matched 16S rRNA gene sequences in cultures, they accounted for ∼70, 50, and 90% of the total ASV abundance for SAR11, SUP05, and *Njordibacter*, respectively (**Fig. S2**). ASVs matching the SUP05, *Njordibacter* and *Marifrigibacter* cultures were more abundant in the EGC, while those matching the SAR11, SAR116, and OM43 cultures were more abundant in the WSC (**Fig. 2; Fig. S3**). Many unique species match the same ASV, and in some cases unique ASVs match the same species, or even different copies of the 16s rRNA gene in the same genome (**Fig. S4**).

**Fig. 2.**
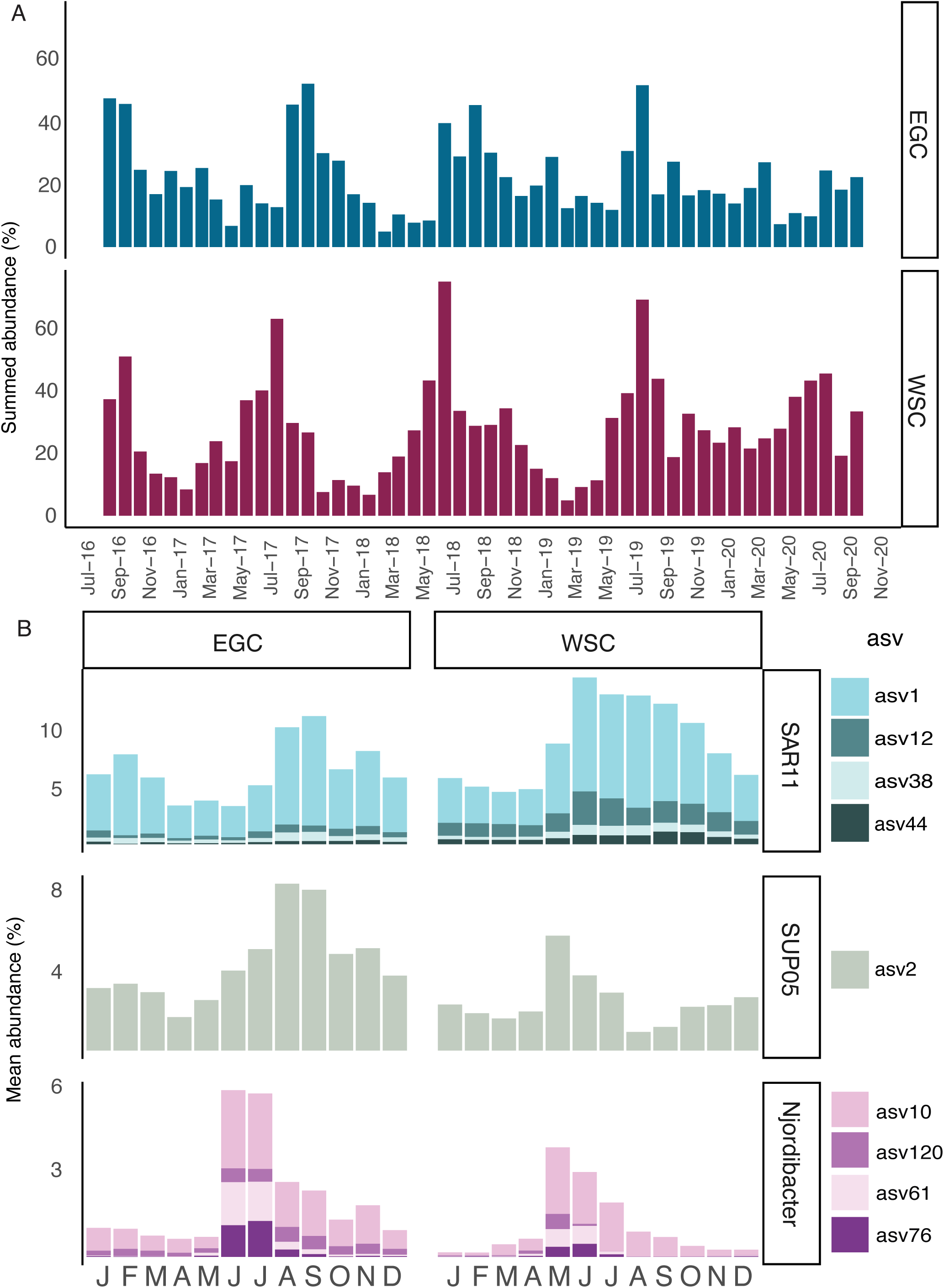
Relative abundance of bacterial ASVs from the Fram Strait time-series with 100% identity to 16S rRNA gene sequences in cultures obtained in this study. **A)** Summed relative abundance of all matching ASVs from August 2016 to September 2020 in the East Greenland Current (EGC) and West Spitsbergen Current (WSC). **B)** Mean relative abundance values calculated for ASVs matching sequenced *Pelagibacter* (SAR11), *Pseudothioglobus* (SUP05), and *Njordibacter* cultures in the EGC (left) and WSC (right). Bars in panel B represent monthly averages between August 2016 and September 2020. Individual ASVs for each lineage in panel B are shown on the right.

### Gene gain, loss, and rearrangement in Arctic SAR11

All 16 SAR11 cultures are members of subclade Ia (**Fig. 3**). Twelve have average nucleotide identity (ANI) values below 95% to each other and two pairs have ANI values above 95% to each other, indicating a collection of cultures with fourteen new species and two new subspecies of SAR11 (**Fig. 3A; Table S3**). Population genomic analysis identified 3,596 total gene clusters, 1,661 singleton gene clusters, and 1,037 core gene clusters (**Fig. 3B**). Each genome encodes between 1,385 – 1,477 unique genes. While the number of shared genes between any two genomes ranges from 1,126 – 1,234, the number of singletons in each genome ranges from 50 – 150.

**Fig. 3.**
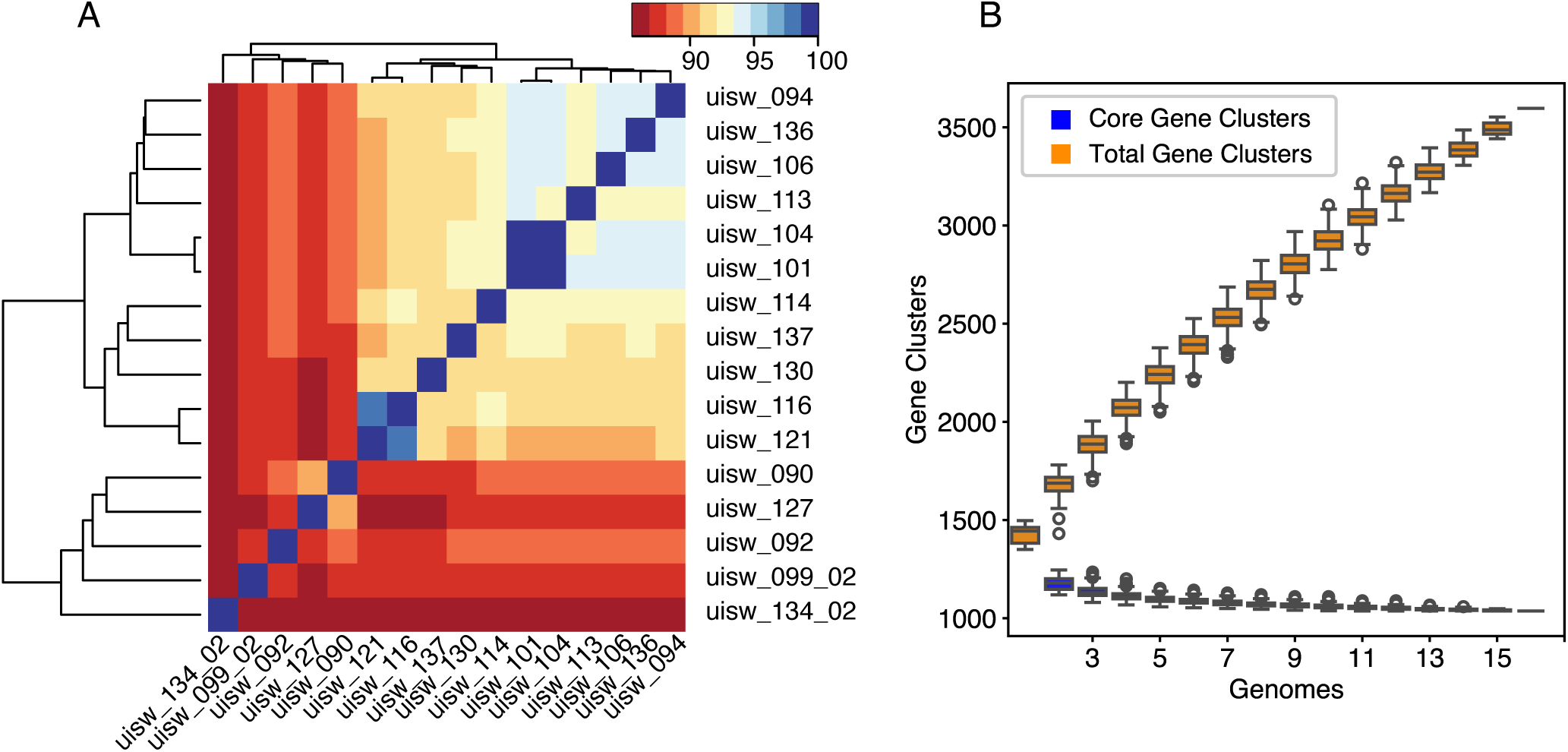
Genomic diversity of Arctic SAR11 cultures. **A**) Clustered average nucleotide identity (ANI) for each pairwise genome comparison. **B**) Gene cluster accumulation curves for the Arctic SAR11 population, total gene clusters (orange) and core gene clusters (blue). Accumulation curves were calculated using 1000 iterations. Analyses were obtained by removing gene cluster paralogs from the anvi summary output file and then converting the data into a presence/absence matrix.

A whole genome alignment was produced between the 16 Arctic SAR11 genomes to identify patterns of gene gain, loss, and rearrangement. The alignment revealed notable patterns of rearrangement (**Fig. 4**). Most notably, there is an 880 Kb section bound by the 23S and 5S rRNA genes in uisw_092 (**Fig. 4A**). Patterns of gene gain, loss, and rearrangement in this region were used to identify a genomic island that has 17-20 genes and is present in half of the sequenced genomes (**Fig. 4B; Table S4**). Genes on the island code for a glycine betaine ABC transport system (ABC trans.), a choline dehydrogenase (CHDH), an aldehyde dehydrogenase (ALDH), a carnitine dehydrogenase (CDH), a Gamma-butyrobetaine dioxygenase (BBOX), and a beta-keto acid cleavage enzyme (BKACE). Gene order in the island is preserved and always upstream of genomic betaine/choline/proline ABC transport genes. Phylogenetic trees constructed using concatenated protein sequences from the island and single-copy core genes are incongruent (**Fig. 4C**), which suggest that the genomic island is acquired by horizontal gene transfer (HGT). Of the 17 island genes, 15 are unique to the island, without identified homologs elsewhere in the genomes, according to gene cluster analysis. The remaining two form distinct phylogenetic clusters from their genomic counterparts, as shown for aldehyde and choline dehydrogenases (**Fig. S5A and S5B**). Phylogenetic analyses of individual island genes indicate that 13 of 17 phylogenies are congruent with each other and with the phylogeny produced using the concatenated island gene set, as shown for aldehyde and choline dehydrogenases (**Fig. S5C and S5D**).

**Fig. 4.**
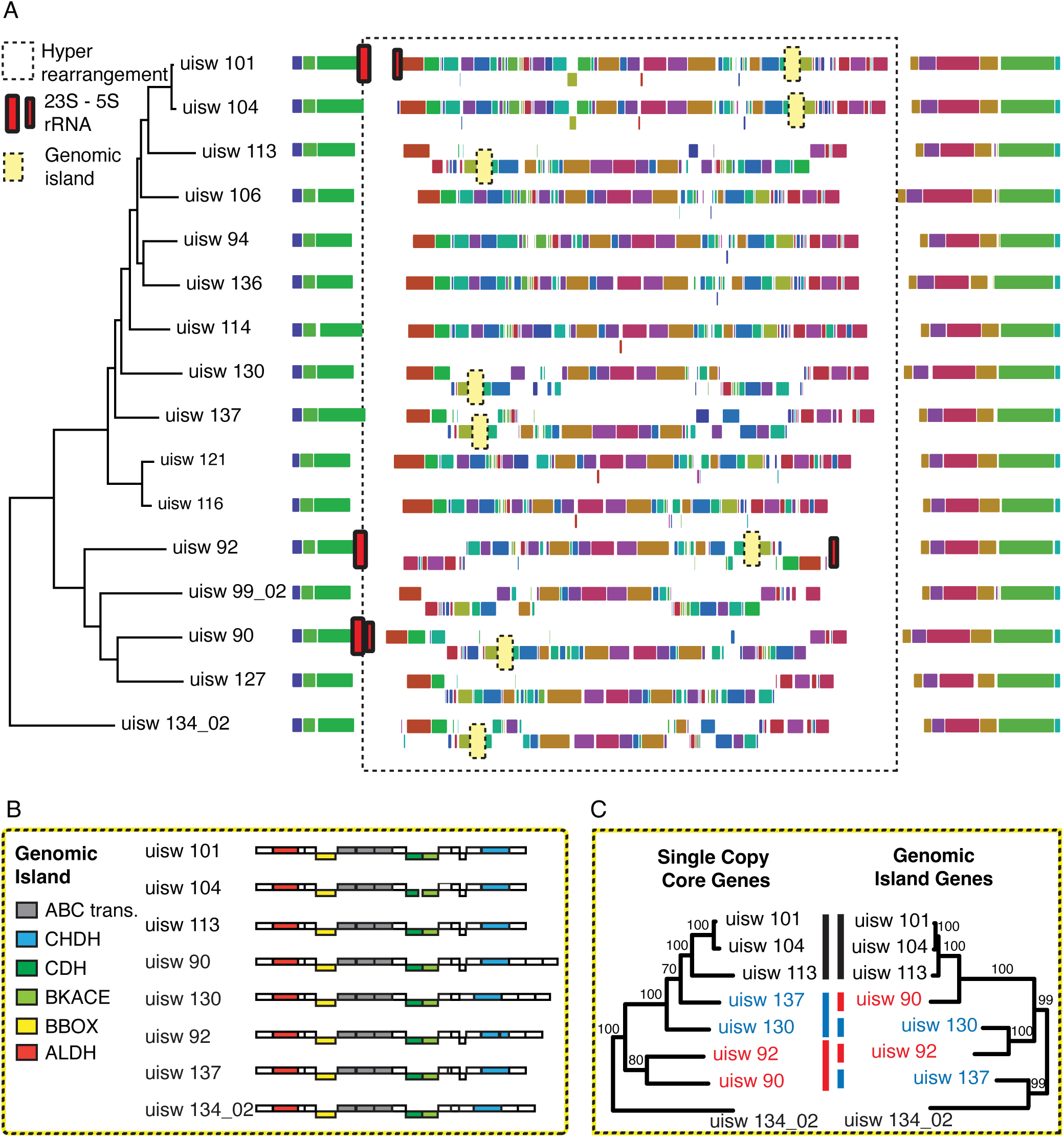
Structure of Arctic SAR11 genomes. **A**) Phylogenomic analysis and whole-genome alignment of Arctic SAR11. **B**) Alignment and gene content of a genomic island identified in eight SAR11 genomes. **C**) Comparison of phylogenetic trees constructed from concatenated protein sequences for single-copy core genes (left) and genomic island genes (right). Phylogenetic trees were constructed with translated and concatenated sequences. Whole genome alignments start with DnaA. The relative position of the 23S and 5S rRNA genes in all genomes is as shown for uisw_101, except for uisw_090 and uisw_092. Enzyme names in panel B are abbreviated as follows: ABC Trans = ABC transporter; CHDH = choline dehydrogenase; CDH = carnitine dehydrogenase; BKACE = beta-keto acid cleavage enzyme; BBOX = gamma-butyrobetaine dioxygenase; ALDH = aldehyde dehydrogenase. Bootstrap values in panel C are displayed at each node. Black, red, and blue color blocks highlight node rearrangements.

We searched for evidence of the Arctic SAR11 genomic island in public databases to see if it was a common but previously unrecognized genomic region. There was no evidence of the island in 18 previously sequenced single-contig SAR11 genomes (**Table S5**). Homologs for genes in the genomic island were identified in environmental databases, most of which were more abundant at polar latitudes (**Fig. 5**). Seven of these genes were rare or absent in samples from lower latitudes, including a lactoylglutathione lyase, a glycerophosphodiester phosphodiesterase, the permease subunit of a proline/glycine betaine ABC transporter, a class II aldolase, a thioesterase, a small multidrug resistance (SMR) transporter, and a drug/metabolite transporter (DMT). Similar but less robust patterns indicating polar distributions were identified for other genes in the genomic island (**Fig. 5**), as has been reported for metagenome read recruitment for SAR11 subclade Ia.1 [75].

**Fig. 5.**
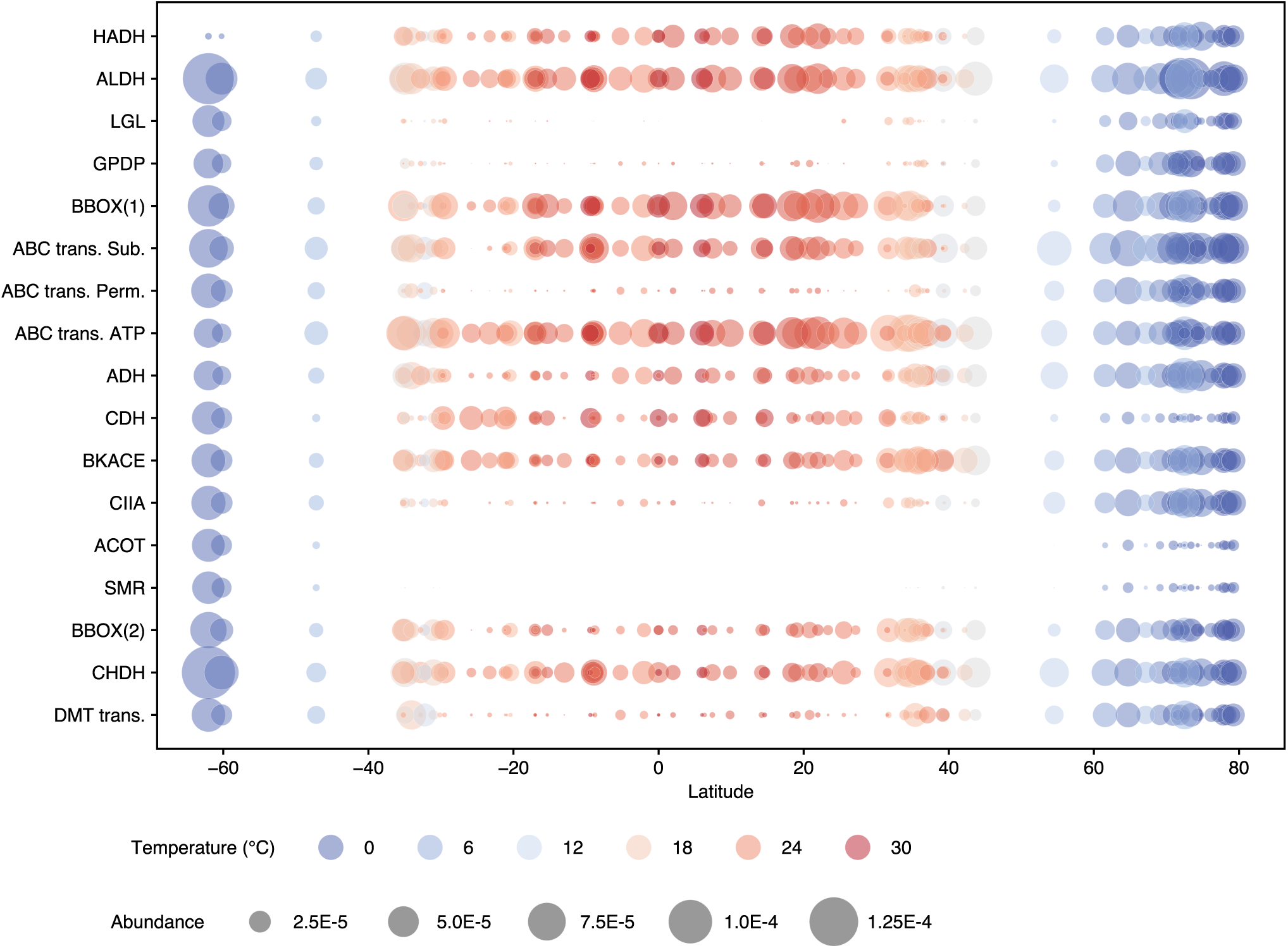
Global distribution of genes identified in the Arctic SAR11 genomic island. Representative nucleotide sequences from uisw_090 were used to identify homologs in the Ocean Gene Atlas OMRGC v2 metaG dataset for all 17 island genes (**Table S4**). Homologs include those with an expect value of E-10 or less. Abundance was normalized to the percent of mapped reads, as previously described [84, 85]. Enzyme names are abbreviated as follows: HADH = 3-hydroxyisobutyrate dehydrogenase; ALDH = aldehyde dehydrogenase; LGL = lactoylglutathione lyase; GPDP = glycerophosphoryl diester phosphodiesterase; BBOX(1) = gamma-butyrobetaine dioxygenase; ABC trans. Sub = ABC transporter substrate binding; ABC trans. Perm. = ABC transporter permease; ABC trans. ATP = ABC transporter ATP binding; ADH = alcohol dehydrogenase; CDH = carnitine dehydrogenase; BKACE = beta-keto acid cleavage enzyme; CIIA = Class II aldolase; ACOT = acyl-CoA thioesterase; SMR = small multidrug resistance transporter; BBOX(2) = gamma-butyrobetaine dioxygenase; CHDH = choline dehydrogenase; DMT trans. = drug/metabolite transporter.

### A new family of Pseudomonadales

A related group of *Pseudomonadales* (*Njordibacter*, named herein) was frequently identified in culture (n=10). Four isolates were selected for whole genome sequencing and used to identify a new family in the order *Pseudomonadales* (**Fig. 6**). The genomes range from 2.4 - 2.9 Mbp in length, have a GC content of 43.8 - 45.1%, and each contain four copies of the 16S rRNA gene. Pairwise ANI values range from 79 - 82% when compared to each other (**Table S2**). One isolate, uisw_056, and a previously characterized MAG from the Arctic [62] have 99.6% ANI. Phylogenomic placement in the Genome Taxonomy Database (GTDB) [76] placed this group in a deeply branching and uncultivated genus of *Nitrincolaceae*, but with low whole genome AAI and 16S rRNA sequence similarity to the most closely related isolate, 60 and 91.7%, respectively. We therefore compared the genomes of 114 isolates in the order *Pseudomonadales* (**Table S6**) to each other and to the 4 *Njordibacter* genomes using AAI and POCP. This yielded over 10,000 pairwise comparisons between 34 genera from 7 families, from which family delineations could be distinguished (**Fig. 6**). Values of 60% AAI and 40% POCP with their closest *Nitrincolaceae* relatives are below the family delineation line, indicating that these *Njordibacter* genomes represents a new family in the order *Pseudomonadales*.

**Fig. 6.**
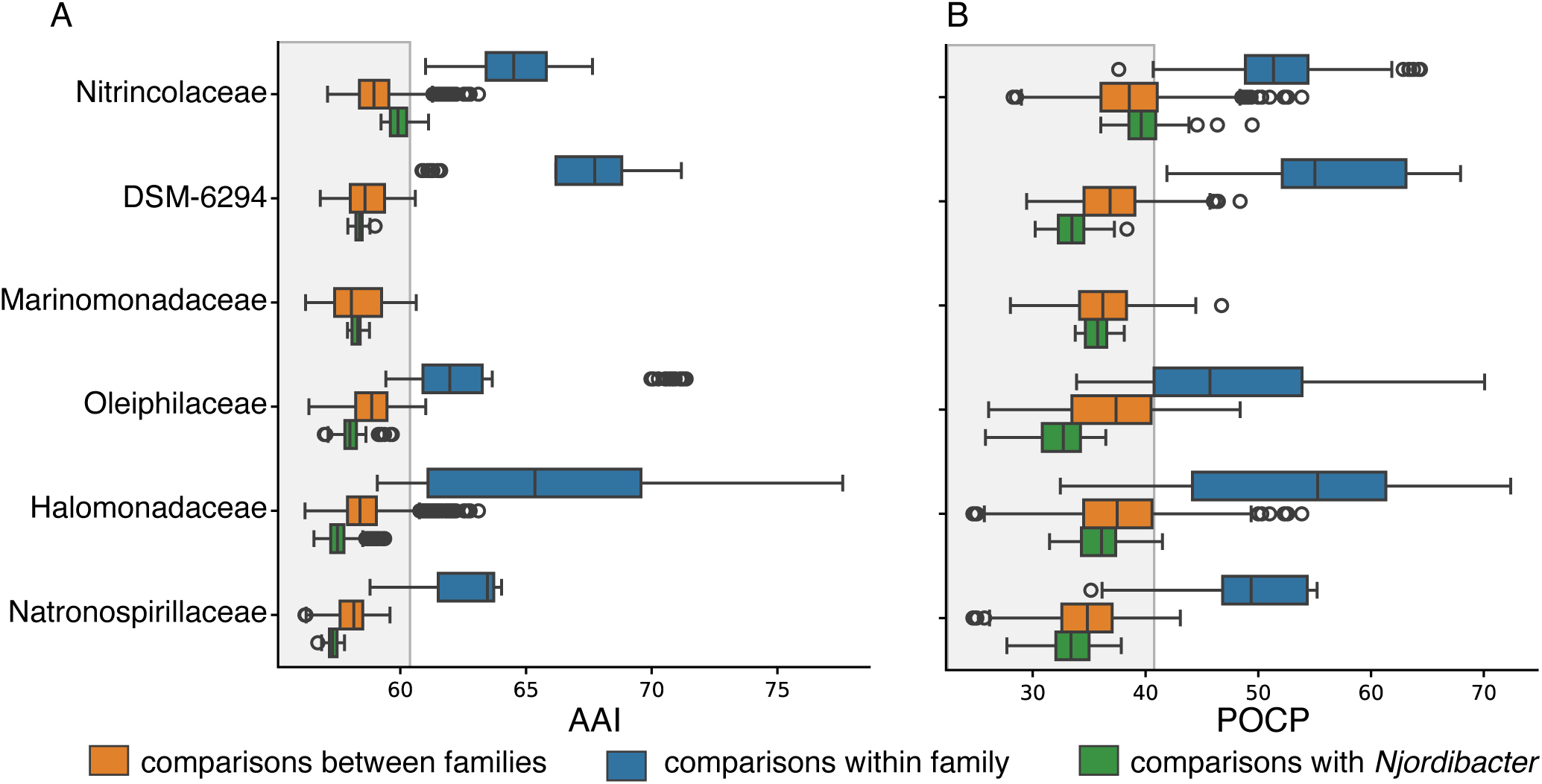
Whole-genome based family and genus delineations in the order *Pseudomonadales*. **A**) All pairwise average amino acid identity (AAI) comparisons. **B**) All pairwise percent of conserved protein (POCP) comparisons. Representatives from the six most closely related families to *Njordibacter* were analyzed (**Table S6**). Circles represent values outside the 1.5 inter-quartile range from the first and third quartile.

## Discussion

Early efforts to assemble genomes of marine bacteria directly from seawater samples produced only partial sequences for the most abundant lineages [77]. While advances in sequencing technology have improved our ability to capture the extensive genetic variation that exists in nature, the extent of genetic diversity within populations remains uncertain, particularly for SAR11 [10, 14]. Our cultivation approach to population genomics provided 34 closed bacterial genomes, including 16 unique SAR11 genomes, from which diversity within a population can be more accurately assessed.

Our analysis identified 1,037 core genes in 16 Arctic SAR11 subclade Ia genomes obtained from just 30 µL of seawater (**Fig. 3**). A similar estimate of 1,047 core genes was obtained in an earlier pangenome analysis of five complete SAR11 subclade Ia genomes isolated from diverse locations in the Atlantic and Pacific Oceans [40].

Similarity between the core genome estimates using 16 genomes from 30 µL of Arctic seawater and five genomes from different oceans highlights the core genome conservation within the SAR11 clade [34, 40], and agrees with a previous metagenomic study, which found greater SAR11 microdiversity between depths of the same location than at the same depths across the global ocean [78]. The SAR11 pangenome analysis also identified 1,962 total unique gene clusters in 5 genomes [40]. The mean number of unique gene clusters in 5 Arctic SAR11 genomes was 2241± 117 (95% CI). This suggests that there is even more genetic diversity in Arctic SAR11 populations. This could be due to the mixing of different populations in Arctic and Atlantic waters [19, 20], or due to enhanced genetic diversity within Arctic populations. For example, Fuhrman and colleagues found that microbial diversity increases with latitude and decreasing temperature [79]. Regardless, our analyses highlight the extent of SAR11 diversity in the Arctic and suggest that additional sequencing is needed to estimate the total number of SAR11 species and subspecies in Arctic populations.

Small gene indels were identified throughout the SAR11 genomes, although most unique genes are co-located in a large ∼50 Kb HVR (HVR2), which is bound by the 23S and 5S rRNA genes [45]. We identified two SAR11 genomes with anomalous gene content in HVR2. Uisw_090 has a single gene and uisw_092 has 995 genes in HRV2 (**Fig. 4A**). The translocation of uisw_092 was used to define a genomic region with a higher frequency of inversions and gene indels relative to regions flanking the origin of replication, which suggests that there is more selective pressure to conserve gene order near the origin of replication.

We identified a genomic island within this 880 Kbp rearrangement region that encodes genes for the uptake and production of glycine betaine (**Fig. 4; Table S4**). Incongruent phylogenies between single-copy core genes and island genes suggests the island is acquired through HGT, and conserved phylogenies for each island gene suggests they are transferred together (**Fig. 4C**; **Fig. S5**). Notably, the island and flanking sequences lack HGT facilitation genes, raising questions about the mechanism of transfer. Homologous recombination has a widespread occurrence between even distantly related SAR11 [46] and has been proposed as a mechanism for the transfer of genomic islands [47]. Homologous recombination has also been proposed as a mechanism for genomic diversification in two other marine clades, HIMB59 (a sister clade to SAR11) and OM43 [80, 81].

Molina-Pardines and colleagues found that a genomic island in HIMB59 had more phosphorus acquisition genes when phosphorus was limiting [80]. A genomic island with genes to use the compatible solute glycine betaine could enhance the survival and metabolic activities of Arctic SAR11 during freezing, as reported for other bacteria [21–23], and may serve as an important methyl donor and source of glycine [24]. Evidence that homologs are absent in more temperate regions and more abundant at high and low latitudes and low temperatures, suggests that the island is an adaption to Arctic conditions (**Fig. 5**). This could give SAR11 cells with the genomic island an advantage during the annual formation and melting of sea-ice.

There are observable differences in the genomic diversity for the two most frequently cultured bacteria in our culture collection, SAR11 and SUP05. Fourteen of the Arctic SAR11 isolates are different species, with ANI values that are <95% to each other (**Fig. 3A**). By comparison, all Arctic SUP05 isolates are subspecies with ANI values that are >95% to each other. Time-series ASV data from the Fram Strait suggest that these are among the most abundant Arctic taxa, representing ∼15% and ∼10% of the community at peak abundance, respectively (**Fig. 2B**). ASV and genomic data both indicate that the SAR11 population is more diverse than the SUP05 population. This suggests that different evolutionary mechanisms are used to maintain diversity in these two lineages. Higher frequencies of genomic rearrangements have been reported as a mechanism for diversification in marine nitrogen fixers [82]. A similar process may help maintain diversity in SUP05 relative to SAR11, although additional closed genomes are needed to quantify differences in the frequencies of gene gain, loss, and rearrangement in both SAR11 and SUP05 populations.

## Conclusion

This cultivation-based population genomics study brought several new families, genera, and species into isolation, and provided new information about the structure of their genomes and adaptations to the Arctic. Half the SAR11 isolates obtained in this study have a genomic island with the potential to enhance the use of glycine betaine, which is an osmolyte produced by phytoplankton throughout the oceans, but which fluctuates annually in cells in polar waters [83]. Our data suggest that adaptation to this fluctuating resource has contributed to the diversification of Arctic SAR11 populations. Several isolates and genomes of other Arctic residents were also identified. *Njordibacter* were frequently identified in our cultured collection, and access to their complete genomes improved family level taxonomic classifications in the order *Pseudomonadales* (**Fig. 6**). Additional cultures and complete genome sequences will improve our understanding of the evolutionary and environmental factors that shape diversity within Arctic populations and elsewhere in the oceans.

## Protologue

*Candidatus* Njordibacter *gen. nov*.

*Njordibacter* (Njord.i.bac’ter. N.L. masc. n. njordi from the Norse word Njord, the god of wind and seas).

*Candidatus* Njordibacteraceae *fam. nov*.

*Njordibacteraceae* (Njord.i.bac’ter.a.ce’ae. N.L. masc. n. *Njordibacter*, type genus of the family; suff. -aceae, ending to denote a family; N.L. fem. pl. n. *Njordibacteraceae*, the family of the genus *Njordibacter*). Description of the family *Njordibacteraceae* is the same as for the genus *Njordibacter*.

*Candidatus* Levibacter *gen. nov*.

*Levibacter* (Levi.bac’ter. N.L. masc. n. levi from the Latin levis, which can mean light in weight. Alluding to the low GC clade of the order *Puniceispirillales*).

*Candidatus* Levibacteraceae *fam. nov*.

*Levibacteraceae* (Levi.bac’ter.a.ce’ae N.L. masc. n. *Levibacter*, type genus of the family; suff. -aceae, ending to denote a family; N.L. fem. pl. n. *Levibacteraceae*, the family of the genus *Levibacter*). The description of the family *Levibacteraceae* is the same as for the genus *Levibacter*.

*Candidatus Ponderosibacter gen. nov*.

*Ponderosibacter* (ponderosi.bac’ter N.L. masc. n. ponderosi from the Latin ponderosus, which can mean heavy or weighty, alluding to the high GC lineage of the order *Puniceispirillales)*.

*Candidatus Marifrigibacter gen. nov*.

*Marifrigibacter* (mari.frigi.bac’ter N.L. masc. n. mari from the Latin mare which means sea and frigi from the Latin frigus which means cold).

## Acknowledgements

Field sampling was supported by the Research Council of Norway (325405), and the Norwegian Institute of Marine Research (Cruise 2023007008). We would like to thank Karley Campbell and Polona Itkin for providing sampling opportunities through the BREATHE sea ice field school, the crew of the RV Kronprins Haakon, and all members of the University of Washington Center for Environmental Genomics for endless support and discussion. We thank everyone in the FRAM consortium for their invaluable role in Arctic long-term observations.

## Data Availability

All data are publicly available at the NCBI under Bioproject PRJNA1129510. Individual genome accession and sequence read archive accession numbers are available in Table S2.

## Author Contributions

MCS performed sampling, cultivation, sequencing, genome analyses, and manuscript writing. MW performed ASV analysis and manuscript editing. SM provided sequencing expertise and manuscript editing. RMM performed genome analyses, manuscript writing, manuscript editing, and project supervision.

## Funding

This work was funded through a National Science Foundation award to RMM (2201310) and a Hall Conservation Genetic Research award to MCS. MW was supported by the SPP 1158 of the German Research Foundation, grant 522416631. SM was supported by the Japan Society for the Promotion of Science through Overseas Research Fellowships.

---

Supplementary Fig. 1. Quality of Oxford Nanopore Technology (ONT)-only whole genome assemblies. ONT-only assembled genomes were compared to high-quality Illumina-ONT hybrid assembled genomes for SAR11 (strain NP1) and SUP05 (strain EF1), with varying levels of sequence coverage. **A**) Mismatches, **B**) indels, and **C**) excess CDS’ identified in ONT-only assemblies relative to hybrid assemblies.

Supplementary Fig. 2. Relative abundance of matching ASVs by lineage. ASVs from the Fram Strait time-series dataset with 100% sequence identity to the 16S rRNA gene sequence of a culture are classified as “matching”, while all others are “non-matching.” Numbers indicate the number of matching and non-matching ASVs for each lineage.

Supplementary Fig. 3. Mean relative abundance values calculated for ASVs with 100% identity to 16S rRNA gene sequences matching *Haliglobus*, *Aquiluna*, SAR116, OM43, *Marifrigibacter*, *Porticoccus*, and *Patiriisocius* in the EGC (left) and WSC (right). Bars represent monthly averages between August 2016 and September 2020. Individual ASVs for each lineage are shown on the right.

Supplementary Fig. 4. Mean relative abundance values calculated for ASVs with 100% identity to 16S rRNA gene sequences matching *Pelagibacter*, *Njordibacter*, and *Pseudothioglobus* in the EGC (left) and WSC (right). Bars represent monthly averages between August 2016 and September 2020. Individual ASVs for each lineage are shown on the right.

Supplementary Fig. 5. Phylogenetic analysis of representative genes on the genomic island **A**) aldehyde dehydrogenase, **B**) choline dehydrogenase, **C**) island aldehyde dehydrogenase rooted with uisw_134_02, and **D**) island choline dehydrogenase rooted with uisw_134_02. Similar genes identified by sequence similarity or annotation are included. Gene originating from the genomic islands are boxed. Bootstrap values over 70 are displayed at the node.

## Notes

### Competing Interest Statement

The authors have declared no competing interest.

## References

1. Rappé MS, Giovannoni SJ. The uncultured microbial majority. Annu Rev Microbiol 2003; 57: 369–394.

2. Lloyd KG, Steen AD, Ladau J, Yin J, Crosby L. Phylogenetically novel uncultured microbial cells dominate earth microbiomes. mSystems 2018; 3: e00055–18.

3. Rappé MS, Connon SA, Vergin KL, Giovannoni SJ. Cultivation of the ubiquitous SAR11 marine bacterioplankton clade. Nature 2002; 418: 630–633.

4. Oh HM, Kang I, Lee K, Jang Y, Lim SI, Cho JC. Complete genome sequence of strain IMCC9063, belonging to SAR11 subgroup 3, isolated from the Arctic Ocean. J Bacteriol 2011; 193: 3379–3380.

5. Oh HM, Kwon KK, Kang I, Kang SG, Lee JH, Kim SJ, et al. Complete genome sequence of *Candidatus* Puniceispirillum marinum IMCC1322, a representative of the SAR116 clade in the Alphaproteobacteria. J Bacteriol 2010; 192: 3240– 3241.

6. Marshall KT, Morris RM. Genome sequence of *Candidatus* Thioglobus singularis strain PS1, a mixotroph from the SUP05 clade of marine Gammaproteobacteria. Genome Announc 2015; 3: e01155–15.

7. Shah V, Chang BX, Morris RM. Cultivation of a chemoautotroph from the SUP05 clade of marine bacteria that produces nitrite and consumes ammonium. ISME J 2016; 11: 263–271.

8. Iverson V, Morris RM, Frazar CD, Berthiaume CT, Morales RL, Armbrust EV. Untangling genomes from metagenomes: revealing an uncultured class of marine Euryarchaeota. Science 2012; 335: 587–590.

9. Walsh DA, Zaikova E, Howes CG, Song YC, Wright JJ, Tringe SG, et al. Metagenome of a versatile chemolithoautotroph from expanding oceanic dead zones. Science 2009; 326: 578–582.

10. Tully BJ, Graham ED, Heidelberg JF. The reconstruction of 2,631 draft metagenome-assembled genomes from the global oceans. Sci Data 2018; 5: 170203.

11. Stepanauskas R, Sieracki ME. Matching phylogeny and metabolism in the uncultured marine bacteria, one cell at a time. Proc Natl Acad Sci U S A 2007; 104: 9052–9057.

12. Stepanauskas R, Fergusson EA, Brown J, Poulton NJ, Tupper B, Labonté JM, et al. Improved genome recovery and integrated cell-size analyses of individual uncultured microbial cells and viral particles. Nat Commun 2017; 8: 84.

13. Landry Z, Swan BK, Herndl GJ, Stepanauskas R, Giovannoni SJ. SAR202 genomes from the dark ocean predict pathways for the oxidation of recalcitrant dissolved organic matter. mBio 2017; 8: e00413–17.

14. Chang T, Gavelis GS, Brown JM, Stepanauskas R. Genomic representativeness and chimerism in large collections of SAGs and MAGs of marine prokaryoplankton. Microbiome 2024; 12: 126.

15. Lim Y, Seo JH, Giovannoni SJ, Kang I, Cho JC. Cultivation of marine bacteria of the SAR202 clade. Nat Commun 2023; 14: 5098.

16. Loman NJ, Quick J, Simpson JT. A complete bacterial genome assembled de novo using only nanopore sequencing data. Nat Methods 2015; 12: 733–735.

17. Wick RR, Judd LM, Holt KE. Assembling the perfect bacterial genome using Oxford Nanopore and Illumina sequencing. PLOS Comput Biol 2023; 19: e1010905.

18. Serreze MC, Barrett AP, Slater AG, Woodgate RA, Aagaard K, Lammers RB, et al. The large-scale freshwater cycle of the Arctic. J Geophys Res: Oceans 2006; 111: C08010.

19. Kwok R. Arctic sea ice thickness, volume, and multiyear ice coverage: losses and coupled variability (1958–2018). Environ Res Lett 2018; 13: 105005.

20. Timmermans M, Marshall J. Understanding Arctic Ocean circulation: a review of ocean dynamics in a changing climate. J Geophys Res: Oceans 2020; 125: e2019JC015665.

21. Raymond-Bouchard I, Goordial J, Zolotarov Y, Ronholm J, Stromvik M, Bakermans C, et al. Conserved genomic and amino acid traits of cold adaptation in subzero-growing Arctic permafrost bacteria. FEMS Microbiol Ecol 2018; 94: fiy023.

22. Torstensson A, Young JN, Carlson LT, Ingalls AE, Deming JW. Use of exogenous glycine betaine and its precursor choline as osmoprotectants in Antarctic sea-ice diatoms. J Phycol 2019; 55: 663–675.

23. Ewert M, Deming JW. Bacterial responses to fluctuations and extremes in temperature and brine salinity at the surface of Arctic winter sea ice. FEMS Microbiol Ecol 2014; 89: 476–489.

24. Boysen AK, Durham BP, Kumler W, Key RS, Heal KR, Carlson LT, et al. Glycine betaine uptake and metabolism in marine microbial communities. Environ Microbiol 2022; 24: 2380–2403.

25. Deming JW, Somers LK, Straube WL, Swartz DG, Macdonell MT. Isolation of an obligately barophilic bacterium and description of a new genus, Colwellia gen. nov. Syst Appl Microbiol 1988; 10: 152–160.

26. Gosink JJ, Woese CR, Staley JT. *Polaribacter* gen. nov., with three new species, *P. irgensii* sp. nov., *P. franzmannii* sp. nov. and *P. filamentus* sp. nov., gas-vacuolate polar marine bacteria of the Cytophaga-Flavobacterium-Bacteroides group and reclassification of ‘*Flectobacillus glomeratus*’ as *Polaribacter glomeratus* comb. nov. Int J Syst Evol Microbiol 1998; 48: 223–235.

27. Brinkmeyer R, Knittel K, Jürgens J, Weyland H, Amann R, Helmke E. Diversity and structure of bacterial communities in Arctic versus Antarctic pack ice. Appl Environ Microbiol 2003; 69: 6610–6619.

28. Zeng Y, Zou Y, Grebmeier JM, He J, Zheng T. Culture-independent and -dependent methods to investigate the diversity of planktonic bacteria in the northern Bering Sea. Polar Biol 2012; 35: 117–129.

29. Rantanen M, Karpechko AYu, Lipponen A, Nordling K, Hyvärinen O, Ruosteenoja K, et al. The Arctic has warmed nearly four times faster than the globe since 1979. Commun Earth Environ 2022; 3: 168.

30. Stroeve J, Notz D. Changing state of Arctic sea ice across all seasons. Environ Res Lett 2018; 13: 103001.

31. Polyakov IV, Pnyushkov AV, Alkire MB, Ashik IM, Baumann TM, Carmack EC, et al. Greater role for Atlantic inflows on sea-ice loss in the Eurasian Basin of the Arctic Ocean. Science 2017; 356: 285–291.

32. Jackson VLN, Grevesse T, Kilias ES, Onda DFL, Young KF, Allen MJ, et al. Vulnerability of Arctic Ocean microbial eukaryotes to sea ice loss. Sci Rep 2024; 14: 28896.

33. Morris RM, Rappe MS, Connon SA, Vergin KL, Siebold WA, Carlson CA, et al. SAR11 clade dominates ocean surface bacterioplankton communities. Nature 2002; 420: 806–810.

34. Giovannoni SJ. SAR11 bacteria: the most abundant plankton in the oceans. Annu Rev Mar Sci 2017; 9: 231–255.

35. Bano N, Hollibaugh JT. Phylogenetic composition of bacterioplankton assemblages from the Arctic Ocean. Appl Environ Microbiol 2002; 68: 505–518.

36. Comeau AM, Li WKW, Tremblay JÉ, Carmack EC, Lovejoy C. Arctic Ocean microbial community structure before and after the 2007 record sea ice minimum. PLoS ONE 2011; 6: e27492.

37. Sousa AGG de, Tomasino MP, Duarte P, Fernández-Méndez M, Assmy P, Ribeiro H, et al. Diversity and composition of pelagic prokaryotic and protist communities in a thin Arctic sea-ice regime. Microb Ecol 2019; 78: 388–408.

38. Collins RE, Rocap G, Deming JW. Persistence of bacterial and archaeal communities in sea ice through an Arctic winter. Environ Microbiol 2010; 12: 1828–1841.

39. Carlson CA, Morris R, Parsons R, Treusch AH, Giovannoni SJ, Vergin K. Seasonal dynamics of SAR11 populations in the euphotic and mesopelagic zones of the northwestern Sargasso Sea. ISME J 2009; 3: 283–295.

40. Grote J, Thrash JC, Huggett MJ, Landry ZC, Carini P, Giovannoni SJ, et al. Streamlining and core genome conservation among highly divergent members of the SAR11 clade. mBio 2012; 3: e00252–12.

41. Thrash CJ, Temperton B, Swan BK, Landry ZC, Woyke T, Delong EF, et al. Single-cell enabled comparative genomics of a deep ocean SAR11 bathytype. ISME J 2014; 8: 1440–1451.

42. Vergin KL, Beszteri B, Monier A, Thrash JC, Temperton B, Treusch AH, et al. High-resolution SAR11 ecotype dynamics at the Bermuda Atlantic Time-series Study site by phylogenetic placement of pyrosequences. ISME J 2013; 7: 1322– 1332.

43. Kraemer S, Ramachandran A, Colatriano D, Lovejoy C, Walsh DA. Diversity and biogeography of SAR11 bacteria from the Arctic Ocean. ISME J 2020; 14: 79–90.

44. Giovannoni SJ, Tripp HJ, Givan S, Podar M, Vergin KL, Baptista D, et al. Genome streamlining in a cosmopolitan oceanic bacterium. Science 2005; 309: 1242–1245.

45. Wilhelm LJ, Tripp HJ, Givan SA, Smith DP, Giovannoni SJ. Natural variation in SAR11 marine bacterioplankton genomes inferred from metagenomic data. Biol Direct 2007; 2: 27.

46. López-Pérez M, Haro-Moreno JM, Coutinho FH, Martinez-Garcia M, Rodriguez-Valera F. The evolutionary success of the marine bacterium SAR11 analyzed through a metagenomic perspective. mSystems 2020; 5: e00605–20.

47. López-Pérez M, Martin-Cuadrado AB, Rodriguez-Valera F. Homologous recombination is involved in the diversity of replacement flexible genomic islands in aquatic prokaryotes. Front Genet 2014; 5: 147.

48. Monaghan EA, Freel KC, Rappé MS. Isolation of SAR11 marine bacteria from cryopreserved seawater. mSystems 2020; 5: e00954–20.

49. Bramucci AR, Focardi A, Rinke C, Hugenholtz P, Tyson GW, Seymour JR, et al. Microvolume DNA extraction methods for microscale amplicon and metagenomic studies. ISME Commun 2021; 1: 79.

50. Quast C, Pruesse E, Yilmaz P, Gerken J, Schweer T, Yarza P, et al. The SILVA ribosomal RNA gene database project: improved data processing and web-based tools. Nucleic Acids Res 2012; 41: D590–D596.

51. Glöckner FO, Yilmaz P, Quast C, Gerken J, Beccati A, Ciuprina A, et al. 25 years of serving the community with ribosomal RNA gene reference databases and tools. J Biotechnol 2017; 261: 169–176.

52. Kolmogorov M, Yuan J, Lin Y, Pevzner PA. Assembly of long, error-prone reads using repeat graphs. Nat Biotechnol 2019; 37: 540–546.

53. Community TG, Abueg LAL, Afgan E, Allart O, Awan AH, Bacon WA, et al. The Galaxy platform for accessible, reproducible, and collaborative data analyses: 2024 update. Nucleic Acids Res 2024; 52: W83–W94.

54. Tatusova T, DiCuccio M, Badretdin A, Chetvernin V, Nawrocki EP, Zaslavsky L, et al. NCBI prokaryotic genome annotation pipeline. Nucleic Acids Res 2016; 44: 6614–6624.

55. Li W, O’Neill KR, Haft DH, DiCuccio M, Chetvernin V, Badretdin A, et al. RefSeq: expanding the prokaryotic genome annotation pipeline reach with protein family model curation. Nucleic Acids Res 2020; 49: D1020–D1028.

56. Morris RM, Cain KR, Hvorecny KL, Kollman JM. Lysogenic host–virus interactions in SAR11 marine bacteria. Nat Microbiol 2020; 5: 1011–1015.

57. Morris RM, Mino S. The complete genome sequences of *Thioglobus autotrophicus* strains EF2 and EF3, isolated from an oxycline in Effingham Inlet, British Columbia. Microbiol Resour Announc 2024; 13: e01118–23.

58. Hall M. Rasusa: randomly subsample sequencing reads to a specified coverage. J Open Source Softw 2022; 7: 3941.

59. Md V, Misra S, Li H, Aluru S. Efficient architecture-aware acceleration of BWA-MEM for multicore systems. In: 2019 IEEE International Parallel and Distributed Processing Symposium (IPDPS); 2019: 314–324.

60. Walker BJ, Abeel T, Shea T, Priest M, Abouelliel A, Sakthikumar S, et al. Pilon: an integrated tool for comprehensive microbial variant detection and genome assembly improvement. PLoS ONE 2014; 9: e112963.

61. Mikheenko A, Prjibelski A, Saveliev V, Antipov D, Gurevich A. Versatile genome assembly evaluation with QUAST-LG. Bioinformatics 2018; 34: i142–i150.

62. Priest T, Appen WJ von, Oldenburg E, Popa O, Torres-Valdés S, Bienhold C, et al. Atlantic water influx and sea-ice cover drive taxonomic and functional shifts in Arctic marine bacterial communities. ISME J 2023; 17: 1612–1625.

63. Priest T, Oldenburg E, Popa O, Dede B, Metfies K, Appen WJ von, et al. Seasonal recurrence and modular assembly of an Arctic pelagic marine microbiome. Nat Commun 2025; 16: 1326.

64. Chaumeil PA, Mussig AJ, Hugenholtz P, Parks DH. GTDB-Tk v2: memory friendly classification with the genome taxonomy database. Bioinformatics 2022; 38: 5315–5316.

65. Kim D, Park S, Chun J. Introducing EzAAI: a pipeline for high throughput calculations of prokaryotic average amino acid identity. J Microbiol 2021; 59: 476–480.

66. Hölzer M. POCP-nf: an automatic Nextflow pipeline for calculating the percentage of conserved proteins in bacterial taxonomy. Bioinformatics 2024; 40: btae175.

67. Pritchard L, Glover RH, Humphris S, Elphinstone JG, Toth IK. Genomics and taxonomy in diagnostics for food security: soft-rotting enterobacterial plant pathogens. Anal Methods 2015; 8: 12–24.

68. Darling AE, Mau B, Perna NT. progressiveMauve: multiple genome alignment with gene gain, loss and rearrangement. PLoS ONE 2010; 5: e11147.

69. Edgar RC. MUSCLE: a multiple sequence alignment method with reduced time and space complexity. BMC Bioinform 2004; 5: 113.

70. Stamatakis A. RAxML version 8: a tool for phylogenetic analysis and post-analysis of large phylogenies. Bioinformatics 2014; 30: 1312–1313.

71. Huerta-Cepas J, Serra F, Bork P. ETE 3: reconstruction, analysis, and visualization of phylogenomic data. Mol Biol Evol 2016; 33: 1635–1638.

72. Eren AM, Kiefl E, Shaiber A, Veseli I, Miller SE, Schechter MS, et al. Community-led, integrated, reproducible multi-omics with anvi’o. Nat Microbiol 2021; 6: 3–6.

73. Delmont TO, Eren AM. Linking pangenomes and metagenomes: the *Prochlorococcus* metapangenome. PeerJ 2018; 6: e4320.

74. Roda-Garcia JJ, Haro-Moreno JM, Huschet LA, Rodriguez-Valera F, López-Pérez M. Phylogenomics of SAR116 clade reveals two subclades with different evolutionary trajectories and an important role in the ocean sulfur cycle. mSystems 2021; 6: e00944–21.

75. Delmont TO, Kiefl E, Kilinc O, Esen OC, Uysal I, Rappé MS, et al. Single-amino acid variants reveal evolutionary processes that shape the biogeography of a global SAR11 subclade. Elife 2019; 8: 1–26.

76. Parks DH, Chuvochina M, Rinke C, Mussig AJ, Chaumeil PA, Hugenholtz P. GTDB: an ongoing census of bacterial and archaeal diversity through a phylogenetically consistent, rank normalized and complete genome-based taxonomy. Nucleic Acids Res 2021; 50: D785–D794.

77. Venter JC, Remington K, Heidelberg JF, Halpern AL, Rusch D, Eisen JA, et al. Environmental genome shotgun sequencing of the Sargasso Sea. Science 2004; 304: 66–74.

78. Haro-Moreno JM, Rodriguez-Valera F, Rosselli R, Martinez-Hernandez F, Roda-Garcia JJ, Gomez ML, et al. Ecogenomics of the SAR11 clade. Environ Microbiol 2020; 22: 1748–1763.

79. Fuhrman JA, Steele JA, Hewson I, Schwalbach MS, Brown MV, Green JL, et al. A latitudinal diversity gradient in planktonic marine bacteria. Proc Natl Acad Sci U S A 2008; 105: 7774–7778.

80. Molina-Pardines C, Haro-Moreno JM, López-Pérez M. Phosphate-related genomic islands as drivers of environmental adaptation in the streamlined marine alphaproteobacterial HIMB59. mSystems 2023; 8: e00898–23.

81. Layoun P, López-Pérez M, Haro-Moreno JM, Haber M, Thrash JC, Henson MW, et al. Flexible genomic island conservation across freshwater and marine Methylophilaceae. ISME J 2024; 18: wrad036.

82. Zehr JP, Bench SR, Mondragon EA, McCarren J, DeLong EF. Low genomic diversity in tropical oceanic N2-fixing cyanobacteria. Proc Natl Acad Sci U S A 2007; 104: 17807–17812.

83. Dawson HM, Connors E, Erazo NG, Sacks JS, Mierzejewski V, Rundell SM, et al. Microbial metabolomic responses to changes in temperature and salinity along the western Antarctic Peninsula. ISME J 2023; 17: 2035–2046.

84. Villar E, Vannier T, Vernette C, Lescot M, Cuenca M, Alexandre A, et al. The Ocean Gene Atlas: exploring the biogeography of plankton genes online. Nucleic Acids Res 2018; 46: W289–W295.

85. Vernette C, Lecubin J, Sánchez P, Coordinators TO, Acinas SG, Babin M, et al. The Ocean Gene Atlas v2.0: online exploration of the biogeography and phylogeny of plankton genes. Nucleic Acids Res 2022; 50: W516–W526.

