## Supplementary figures and images for "Genome restructuring and adaptation in Arctic marine bacteria"

### supplemental_figure_1

Fig. S1
Sadler et al.

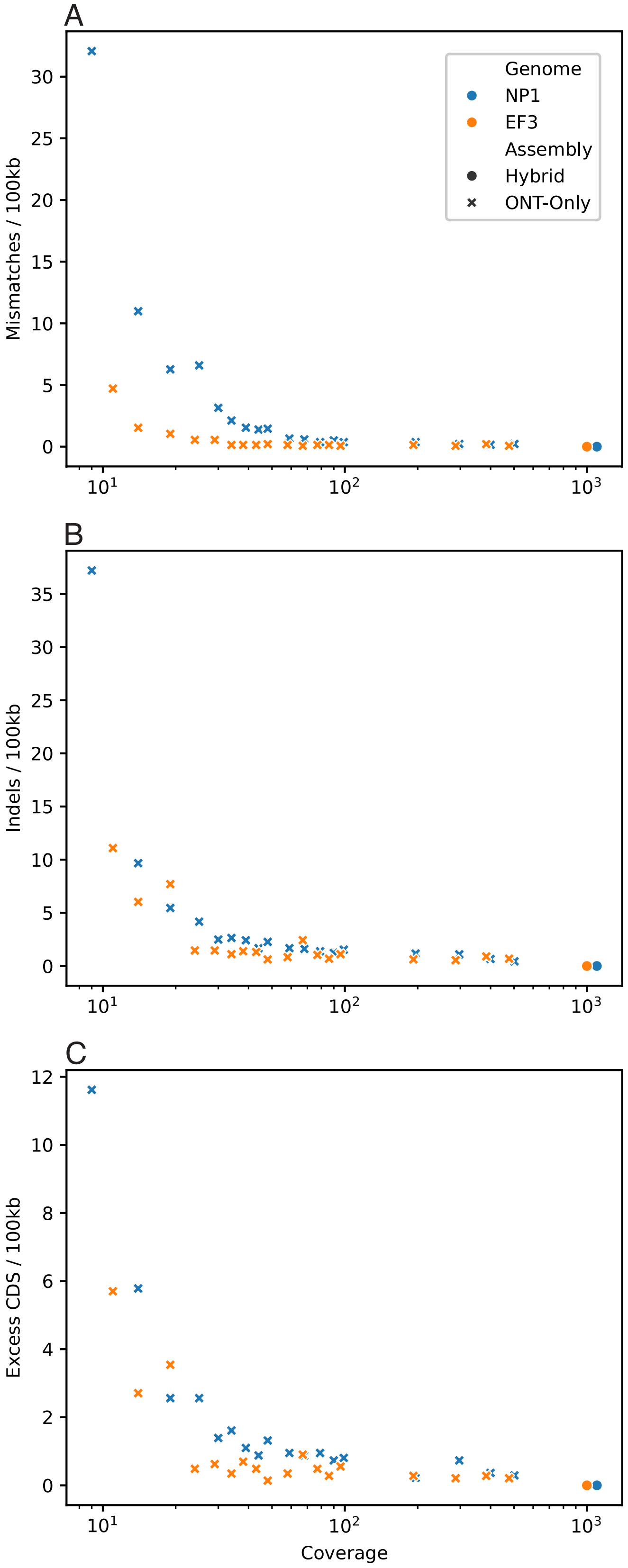

### supplemental_figure_2

Fig. S2
Sadler et al.

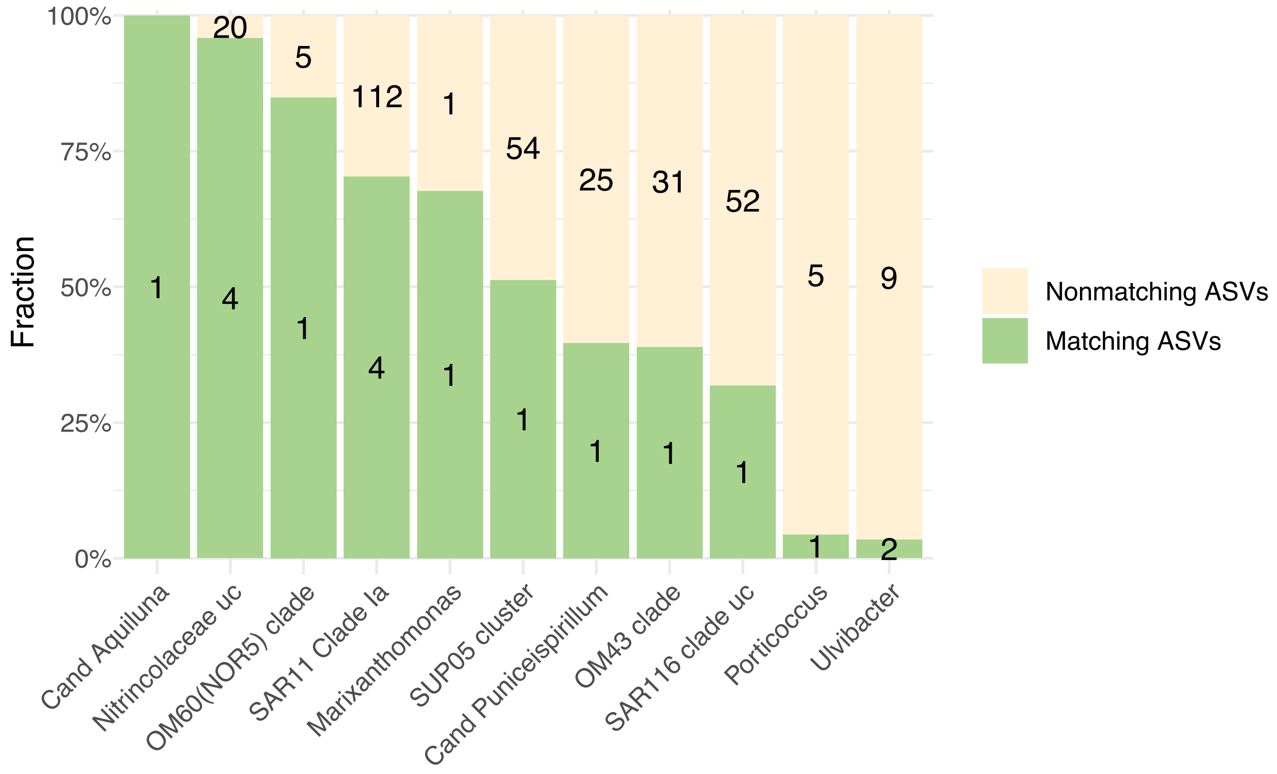

### supplemental_figure_3

Fig. S3
Sadler et al.

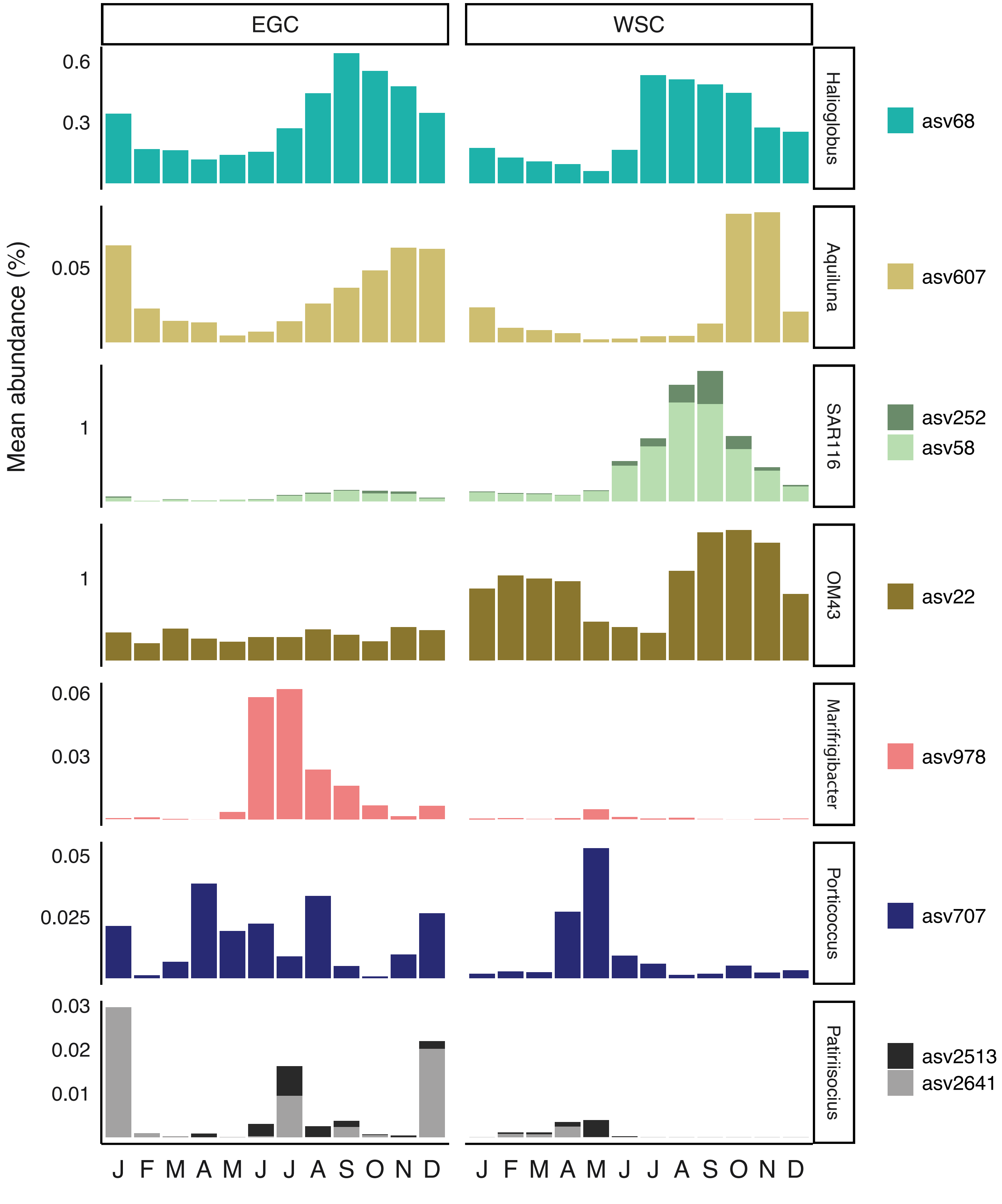

### supplemental_figure_4

Fig. S4
Sadler et al.

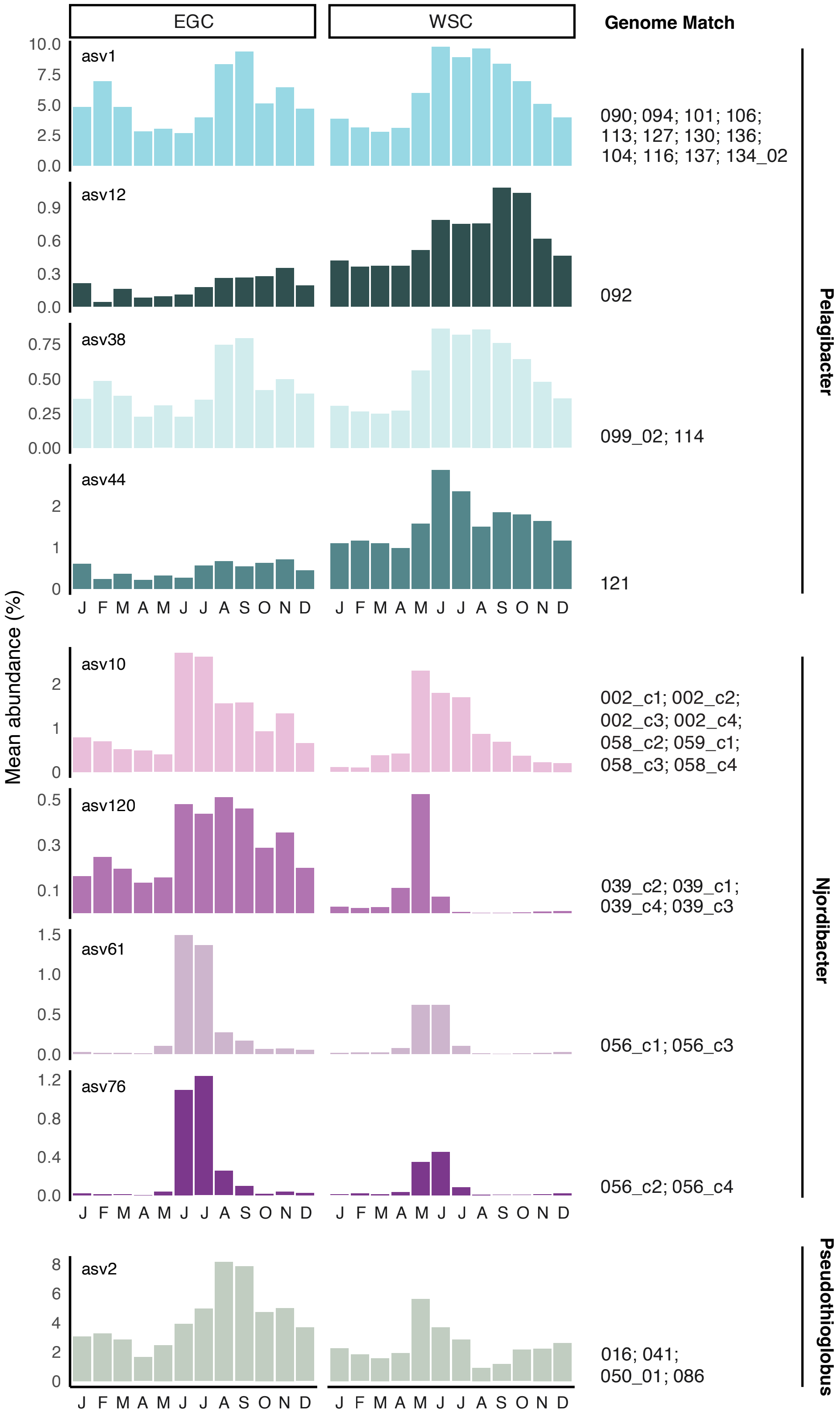

### supplemental_figure_5

Fig. S5
Sadler et al.

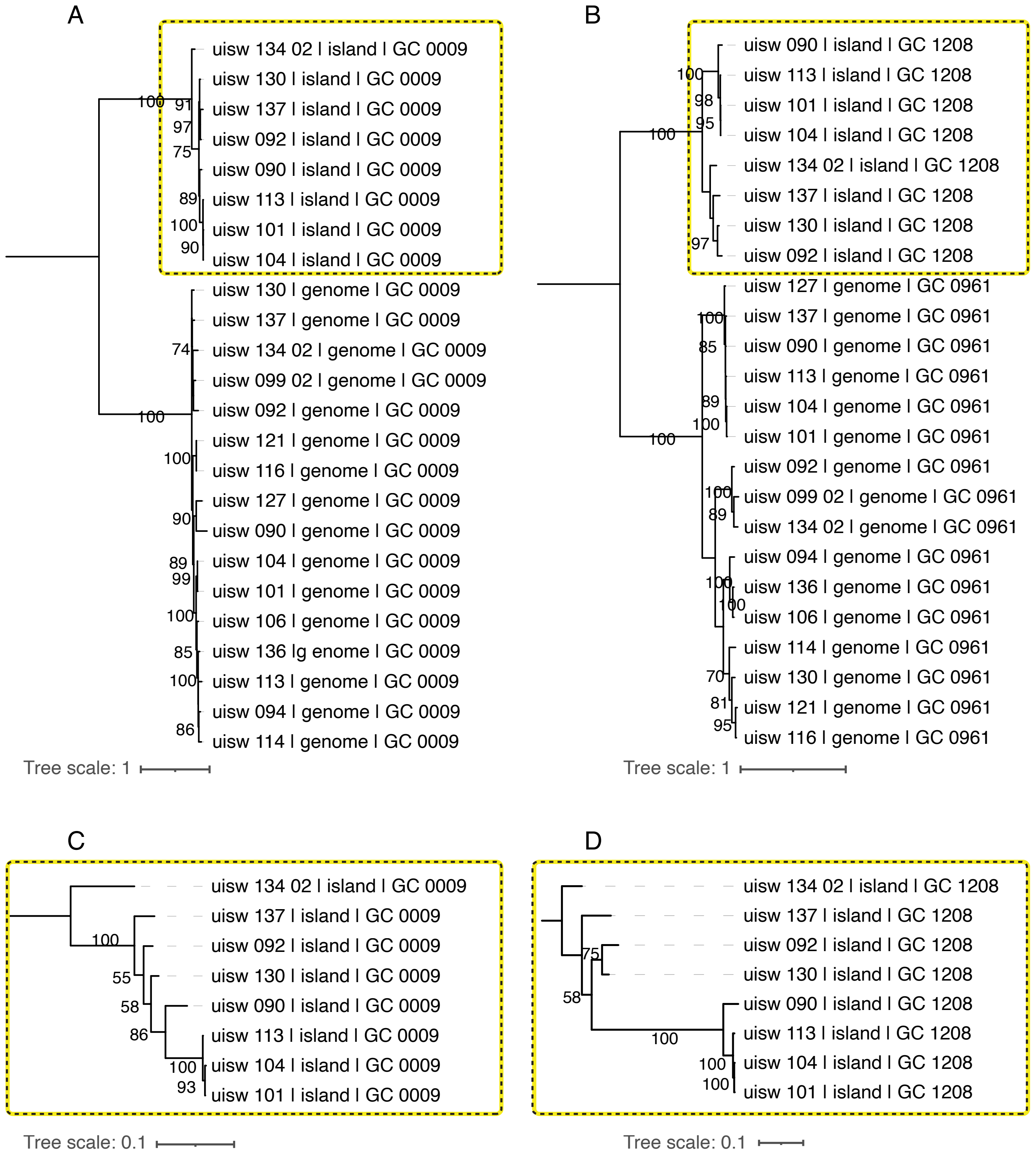
